# Cryo-EM structure of a single-chain β1-adrenoceptor – AmpC β-lactamase fusion protein

**DOI:** 10.1101/2021.09.25.461805

**Authors:** Gabriella Collu, Inayathulla Mohammed, Aleix Lafita, Tobias Bierig, Emiliya Poghosyan, Spencer Bliven, Pavel Afanasyev, Roger M. Benoit

## Abstract

The insertion of fusion proteins has enabled the crystallization of a wide range of G-protein-coupled receptors. Here, we adapted this engineering strategy to cryo-electron microscopy (cryo-EM). We inserted the soluble protein AmpC β-lactamase into the third intracellular loop (ICL3) of ultra-thermostable β1-adrenoceptor (β1AR) via chimeric helix fusions. Biochemical and biophysical characterization showed that the resulting fusion protein after expression, solubilization and purification was monodisperse and able to bind the known β1AR weak partial agonist cyanopindolol, and the antagonist propranolol. The protein particles comprised sufficient mass and discernable structural features to elucidate its cryo-EM structure in complex with cyanopindolol without any natural (G-proteins, arrestins) or artificial (Nanobodies, DARPins) binding partners, to an overall resolution of 4.2 Å. The seven-helix architecture and helix eight, as well as both GPCR - AmpC β-lactamase connections are clearly resolved. β1AR is in its inactive conformation. 3D variability analysis revealed significant flexibility between the two protein domains and within the GPCR helices, offering insights into conformational dynamics. The map contains clear density for the cyanopindolol. The fusion protein geometry theoretically fits a wide range of class A GPCRs, presenting a powerful platform for structure elucidation of a diverse array of class A GPCR – ligand complexes by cryo-EM in the inactive receptor state. The approach furthermore holds potential for structure elucidation of GPCRs in the absence of ligands.

## Introduction

G-protein-coupled receptors (GPCRs) interact with extracellular signaling molecules and transduce the information across the cell membrane through conformational changes within the GPCRs, resulting in various intracellular responses.^1^ Non-rhodopsin GPCRs were for a long time considered among the most challenging protein drug targets for structural studies. The first structures of class A GPCRs were published in 2007^2,3^ and in 2008^4^. Major bottlenecks for the structural biology studies of GPCRs include protein production, for example due to low expression levels or low protein stability, and crystallization, which can be challenging due to high conformational flexibility or a limited area of accessible hydrophilic surface.

Three important strategies that revolutionized GPCR structural studies were the introduction of point mutations to improve the thermostability of the receptors,^4,5^ the binding of specific antigen-binding fragments (Fab),^3^ and the insertion of fusion proteins to stabilize a specific receptor conformation and/or to increase the soluble surface area for promoting the formation of ordered crystal contacts.^2^

More recently, cryo-electron microscopy (cryo-EM) has emerged as a powerful alternative to X-ray crystallography and represents the next major progress in the field of GPCR structural biology. Protein structure elucidation at near-atomic resolution by single particle cryo-EM has seen an impressive increase in the number of structures published through the “resolution revolution”, enabled by important technological advances in specimen preparation, as well as the development of direct electron detectors, improved microscopes and improvements in data analysis.^6^ The major advantages of using cryo-EM for structural studies are that only microgram quantities of protein are needed, crystallization is not required, and different conformational states can be revealed. Nonetheless, due to their relatively small molecular mass (∼45 kDa), combined with the relatively featureless shape within the detergent micelle, GPCRs remain challenging targets for single particles analysis, owing to the low signal-to-noise ratio and to the absence of prominent structural features that would be needed to achieve high accuracy in the determination of the angular orientation for 3D reconstructions, as well as heterogeneity of the protein complexes.^7,8^ For small proteins such as GPCRs, a variety of factors are important to obtain optimal results in cryo-EM.^9^ In 2017, the first GPCR structures were solved by cryo-EM, both belonging to class B: the calcitonin receptor (CTR)^10^ and the glucagon-like peptide-1 receptor (GLP-1)^11^. The structures of these receptors were solved in complex with the heterotrimeric G-proteins, in the Zhang et al.^11^ structure in addition with a specific nanobody. The G-protein complexes had sufficient molecular mass for obtaining a good signal-to-noise ratio in single-particle cryo-EM data collection and sufficiently clearly identifiable structural features for particle alignment. The complex formation relies on the binding of an agonist molecule that activates the receptor, therefore leading to the elucidation of active conformation structures with G-protein or arrestin coupling (e.g. Refs^10–13^). However, this approach is limited by the necessity of agonist-induced conformational changes that facilitate the binding of these partners, making it challenging to obtain high-resolution structures in the absence of such ligands. Therefore, to date, cryo-EM structures of GPCRs in their inactive conformation and apo structures are underrepresented.

To address this challenge, we optimized a molecular engineering strategy widely used in X-ray crystallography to apply it for cryo-EM studies. By searching the Protein Data Bank^14^ (https://www.rcsb.org/) using a BioJava script, we identified AmpC β-lactamase as a relatively large protein (39 kDa) that is well suited for insertion into intracellular loop 3 (ICL3) of class A GPCRs via genetically extended helix fusions.

We designed and cloned single-chain β1AR - AmpC β-lactamase fusion proteins.^15^ Here, we present the cryo-EM structure of a β1AR - AmpC β-lactamase fusion protein in complex with the weak partial agonist cyanopindolol at 4.2 Å resolution.

## Results

### Construct design

To study the structure of cyanopindolol-bound ultra-thermostable β1-adrenergic receptor by cryo-EM, we engineered a single-chain GPCR-fusion protein of sufficient mass and rigidity to allow cryo-EM structural analysis. To identify a suited fusion protein that adds enough mass to the GPCR and that can be connected to the GPCR at ICL3 via rigid, extended α-helices, structures in the PDB were filtered for proteins fulfilling the following criteria: (1) the protein chains are composed of more than 300 residues, (2) N- and C-termini are α-helical, (3) the terminal helices are approximately antiparallel to each other, and (4) the terminal helices are close to 11 Å apart. These constraints were chosen to enable the protein to extend TM5 and TM6 in the ICL3 of class A GPCRs with minimal disruption. Structures from the resulting list of hits were manually inspected, following additional selection criteria: membrane proteins and oligomeric proteins were excluded, and proteins with available high-resolution structures were prioritized.

The AmpC β-lactamase^16^ (PDB:1FCO) from *E. coli* fulfilled all the desired criteria and we therefore used it for insertion into β1AR. The N- and C-terminal helices of β-lactamase were directly fused to the transmembrane helices TM5 and TM6 of β1AR.

Nine different constructs were designed, in which the transition between TM5 of the receptor and the N-terminus of the fusion protein, and the connection between the C-terminus of the fusion protein to TM6 of the receptor, were optimized. The initial construct, which we named construct 0/0, was designed based on visual inspection of the PDB structures of a stabilized turkey β1-adrenergic receptor^4^ (PDB:2VT4) and of the fusion protein^16^ (PDB:1FCO)(Figure 1).

**Figure 1.**
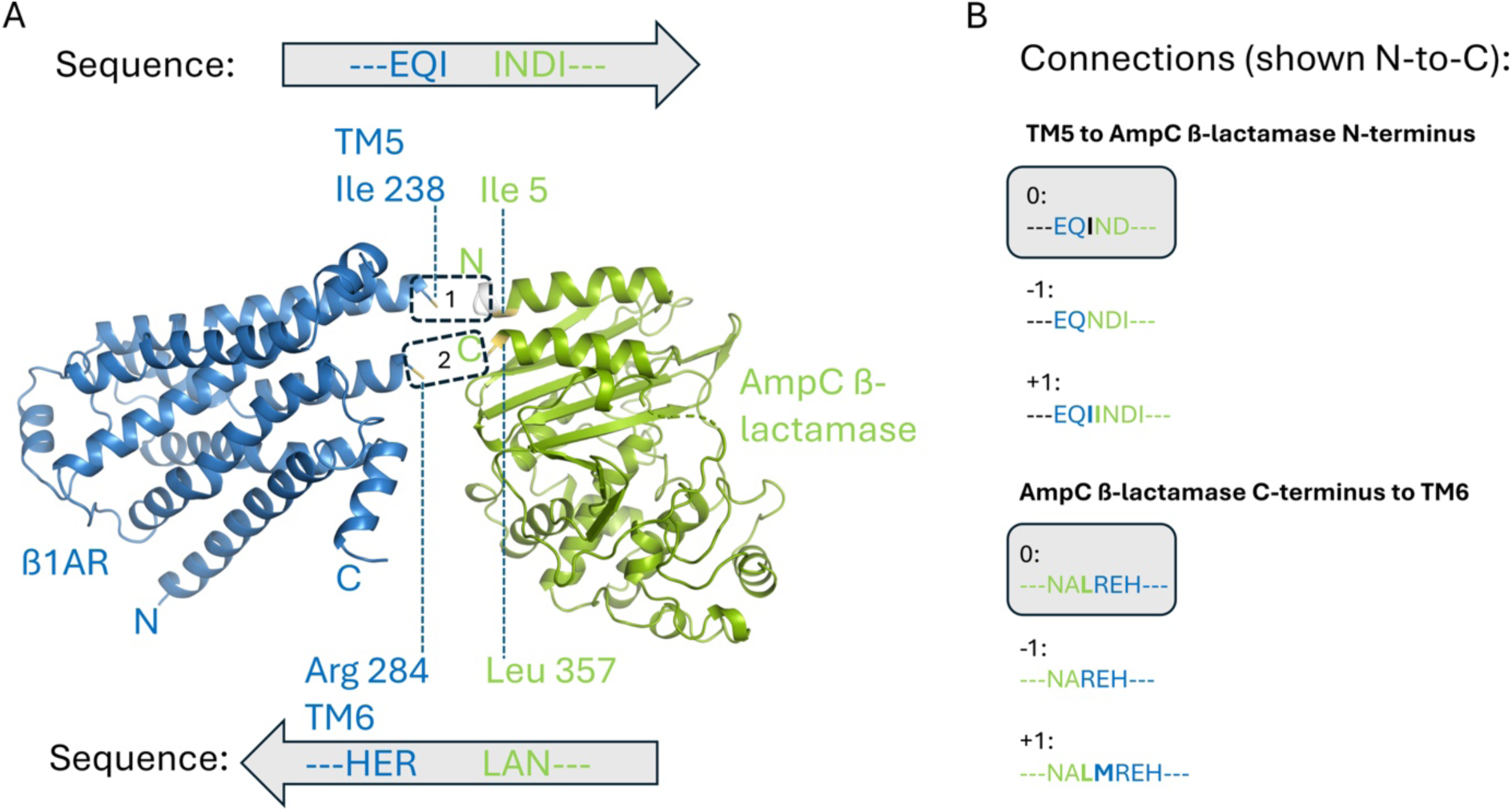
Construct design. **A)** A structure of a stabilized turkey β1-adrenergic receptor^4^ (blue; PDB:2VT4) and a structure of AmpC β-lactamase^16^ (green; PDB:1FCO) were manually aligned to design two chimeric helices that connect the proteins. Chimeric helix 1 connects the C-terminal end of the β1AR transmembrane helix 5 to the N-terminal helix of AmpC β-lactamase (dashed box 1, left-to-right). Chimeric helix 2 connects the C-terminal helix of AmpC β-lactamase to the N-terminal end of β1AR transmembrane helix 6 (dashed box 2, right-to-left). The N- and C-termini of β1AR and AmpC β-lactamase are indicated. **B)** Based on this alignment, initial connections were designed (construct 0/0). For the additional constructs, each of the two connections was either kept at 0 or lengthened or shortened by one amino acid, using only original sequences from β1AR and AmpC β-lactamase, in all possible combinations, resulting in 9 constructs with slightly different linker combinations.

The remaining eight constructs were generated by adding or removing one codon at each of the receptor-fusion protein transitions.^15^ The sequence of our cryo-EM construct was based on the ultra-thermostable mutant β1AR-JM50.^17,18^

The constructs, in a pcDNA 4/TO mammalian expression vector, included a C-terminal flexible linker comprising a human rhinovirus 3C protease (HRV 3C) cleavage site, followed by an eGFP for detection, and a twin Strep-tag for purification.

### Protein expression and purification

The GPCR fusion proteins were produced in stably transfected^19^ HEK293S GnTI(-)^20^ cells. For expression tests, adherent cell cultures were used; for protein production, the cells were grown as suspension cultures.

To extract the protein of interest, membranes were solubilized using n-Dodecyl-β-D-Maltoside (β-DDM). Purification was performed by Strep-Tactin affinity chromatography followed by size-exclusion chromatography (SEC).

The purification of the proteins for cryo-EM analysis was performed in the presence of the high-affinity β1AR-binder cyanopindolol. The purification of proteins for biophysical characterization was performed in the absence of ligands. For cryo-EM analysis and for the biophysical characterization of the constructs, the C-terminal eGFP, which was fused to the protein of interest via a flexible linker, was removed by cleavage with HRV 3C protease (Figure 2 A and B).

**Figure 2.**
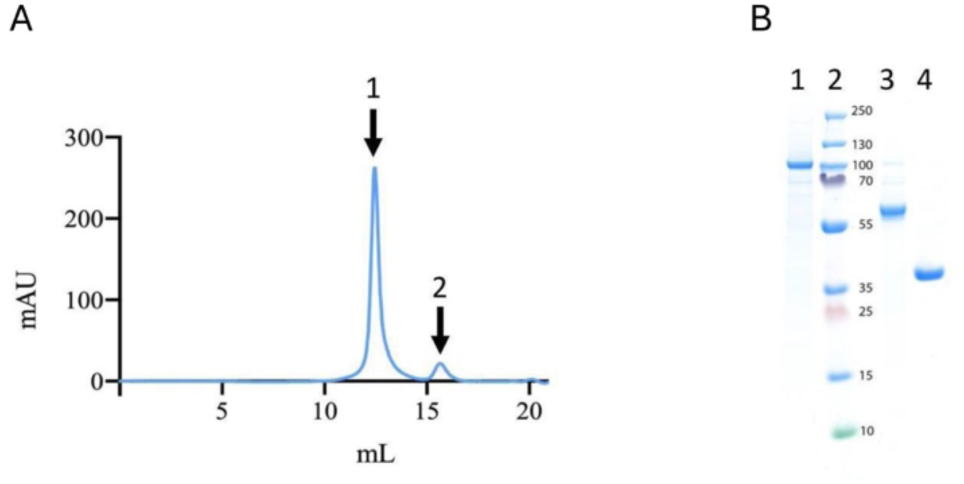
Protein purification. **A)** Size-exclusion chromatography profile of the β1AR-AmpC β-lactamase fusion protein construct 0/0 (peak 1), with the C-terminal eGFP (peak 2) cleaved off using HRV-3C protease. **B)** SDS-PAGE analysis: (1) uncleaved β1AR-AmpC β-lactamase construct with eGFP, (2) PageRuler Plus Prestained Protein Ladder (Thermo Scientific), (3) cleaved β1AR - AmpC β-lactamase construct, and (4), cleaved-off eGFP.

### Ligand-binding assays

N-[4-(7-diethylamino-4-methyl-3-coumarinyl)phenyl]maleimide (CPM) based fluorescent thermal stability^21^ assays were performed on the three most diverse constructs, named −1/-1, 0/0 and +1/+1, to assess the stability and correct folding of the receptor in the fusion protein. The thermostability of the proteins was measured upon addition of increasing concentrations of the weak partial β1AR agonist cyanopindolol, or the β1AR antagonist propranolol. The addition of the small molecule cyanopindolol resulted in a strong stabilizing effect on all tested constructs, while the addition of the small molecule propranolol had a less strong, but still significant stabilizing effect (data not shown). We proceeded with the 0/0 construct for structural studies.

### Structure of the β1AR – AmpC β-lactamase fusion protein

#### Cryo-electron microscopy

We solved the cryo-EM structure of the β1AR fusion protein to an overall resolution of about 4.2 Å. The cryo-EM map revealed the connectivity of all eight helices of the receptor. In contrast to the crystal structure,^18^ where a single conformation is embedded in the crystal lattice, the cryo-EM data reveals conformational heterogeneity within the receptor and between the receptor and the fusion protein. 3D variability analysis shows movements between β1AR and AmpC β-lactamase, and variability in the positions of the α-helices of β1AR with respect to each other (Supplementary Movie 1), revealing a conformational flexibility that is typical for GPCRs. The density for the ligand cyanopindolol, which binds deeply in the center of the extracellular region of the receptor, is defined in the cryo-EM map and allows determination of the overall ligand binding mode. The sharpened map of the overall structure, the real-space refined model and a close-up view of the ligand-binding site are shown in Figure 3.

**Figure 3.**
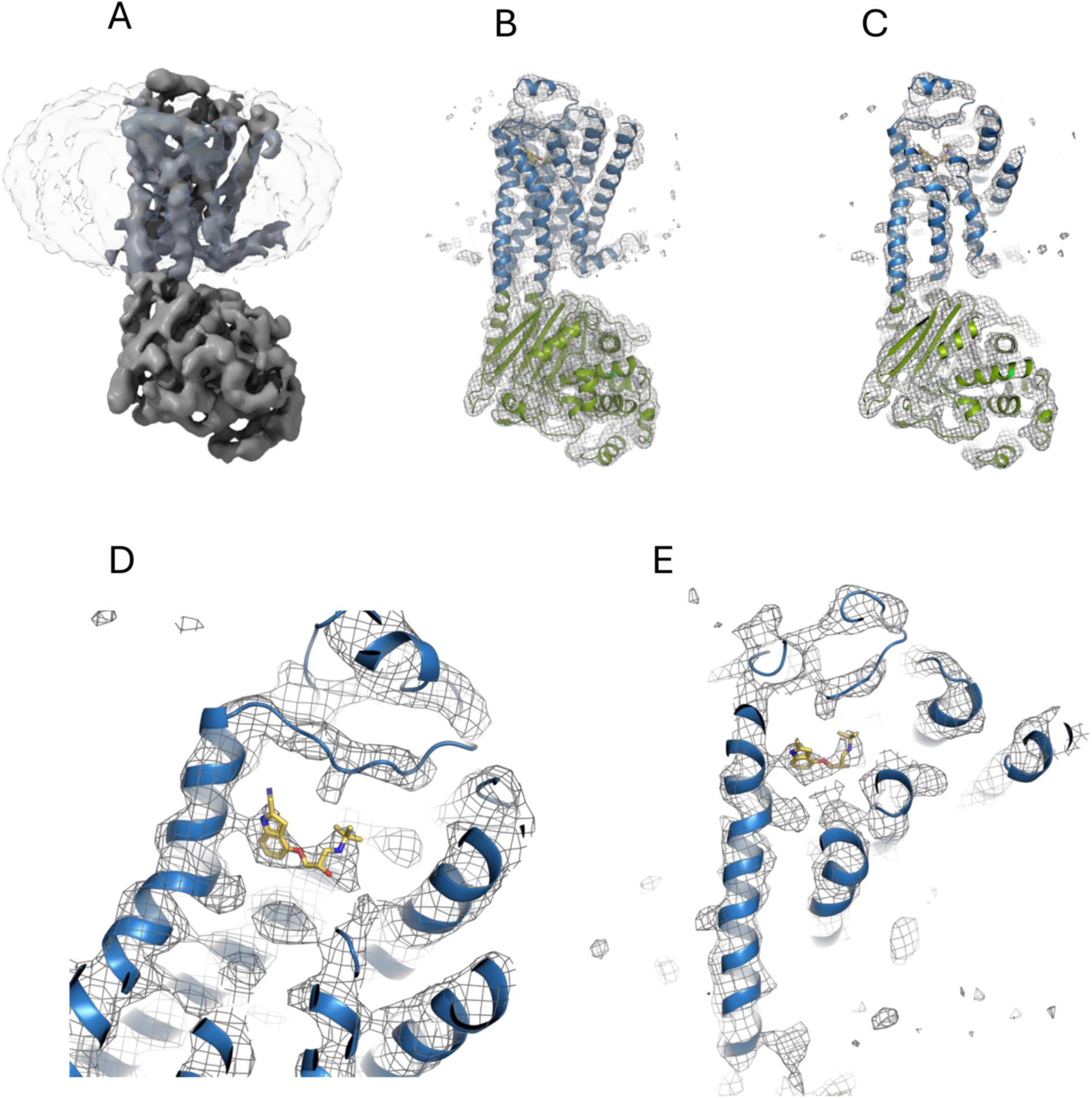
Overall structure and ligand-binding site. **A)** Cryo-EM map of the β1AR fusion protein at 4.2 Å resolution. **B, C)** Real-space refined model (cartoon representation) of the β1-adrenoceptor – AmpC β-lactamase fusion protein fitted in the cryo-EM map, overall (B) and slab (C) view. **D and E)** Close-up slab views of the ligand binding site. In B-E, the β1-adrenoceptor is shown in blue, AmpC β-lactamase is shown in green, and the ligand cyanopindolol is shown in yellow and atom colors (blue = nitrogen, red = oxygen). The Figure was created using ChimeraX^22^ and Pymol^23^.

The resolution was limited by a strong conformational heterogeneity (Supplementary Movie 1) and the number of particles. Locally, particularly at the ligand binding site, the resolution and the quality of the map are significantly better than on the periphery (Supplementary Figure 1, see local resolution), which allowed us to build the atomic model especially at the ligand binding area with strong confidence. The quality of the model was validated with the existing X-ray data.

#### Helix 8 variability

In a physiological environment, helix 8 of adrenergic receptors is anchored to the cytoplasmic face of the cell membrane through palmitoyl groups.^24^ In our construct, the palmitoylation site (C358A) was removed by mutagenesis, and the protein was solubilized in detergent, hence no membrane was present. Despite the lack of membrane anchoring or crystal contacts, helix 8 is defined in the cryo-EM map from Pro 347 up to Arg 361. Due to flexibility, the density is difficult to interpret after Lys 354. Helix 8 adopts a typical position slanted upwards at a shallow angle towards the expected location of a cell membrane. It is likely that the conformation of helix 8 is stabilized by the micelle.

#### Model building and interpretation

An initial model was generated based on an AlphaFold2^25^ model of the fusion protein, and high-resolution crystal structures of both protein components. The model was fitted into the cryo-EM map and refined in iterative cycles of model building and refinement.

#### Ligand binding site

The cyanopindolol binding site was analyzed and validated by comparison to the crystal structure (PDB:4BVN)^18^. In our model, Asp 121^3.32^, Ser 211^5.42^ and Tyr 333^7.53^ are in hydrogen bonding distance to the ligand, in agreement with the crystal structure. Superscripts indicate Ballesteros-Weinstein nomenclature.^26^ Asn 329^7.39^, which makes a hydrogen bond with the ligand in the crystal structure, is slightly more distant from the ligand N2 in the cryo-EM model (3.7 vs. 2.8 Å). Asp 121^3.32^, Asn 310^6.55^ and Tyr333^7.53^ are reasonably well defined in the cryo-EM map, the other hydrogen-bonding residues are located as expected from the overall C_α_-positions. Overall, we could typically build aromatic residues with strong confidence.

The ligand is furthermore lined by residues that make non-polar contacts: Trp 117, Thr 118, Val 122, Val 125, Thr 126, Phe 201, Thr 203, Tyr 207, Ala 208, Ser 212, Ser 215, Trp 303, Phe 306, Phe 307, and Phe 325. Thr 117, Thr 118, Val 125, and Trp 303 are defined in the cryo-EM map.

#### Activation state

Superposition of our cryo-EM structure with the crystal structure of the β1AR-JM50 construct^18^ (PDB:4BVN) shows that, based on the overall conformation of the transmembrane helices, the receptor adopts an inactive (R) conformation, in agreement with the X-ray crystal structure of the same receptor without a fusion protein (Figure 4).

**Figure 4.**
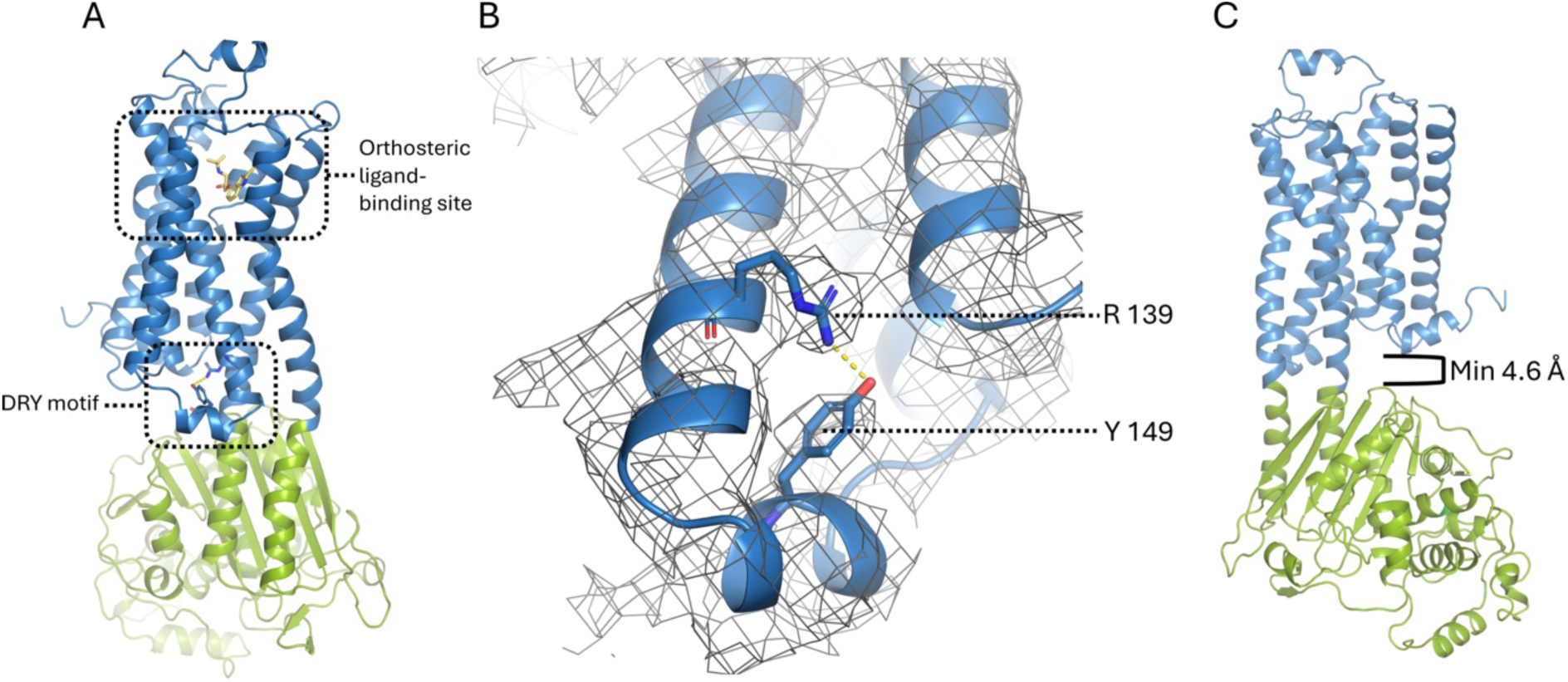
DRY motif and fusion protein geometry. The fusion protein is shown in blue (β1AR) and green (AmpC β-lactamase). **A)** Orthosteric ligand binding site and the DRY motif shown in the context of the overall structure. **B)** Close-up view of the DRY motif. The cryo-EM density map is shown as a mesh. **C)** The minimum distance between the two proteins (except at the covalently linked chimeric helices) is ∼4.6 Å, therefore, steric hindrance that could interfere with inter-helix conformational changes (except between the fused TM5 and 6) is unlikely. The Figure was created using Pymol^23^.

The DRY motif (Figure 4 A and B) in TM α-helix 3 is composed of Asp 138^3.49^, Arg 139^3.50^ and Tyr 140^3.51^, and has previously been shown to interact with Tyr149 from the short α-helix in ICL2^4^ in the inactive state. Arg 139^3.50^ and Tyr 149 are defined in the cryo-EM map and are in agreement with this interaction (Figure 4 B).

The minimal distance between the two proteins is 4.6 Å (Figure 4 C). It is therefore unlikely that the AmpC β-lactamase hinders the conformational freedom of β1AR in regions other than the directly fused helices.

Two disulphide bonds hold extracellular loop 2 (ECL2) in place with respect to TM3, and ECL2 has been suggested to control access to the orthosteric ligand binding site.^27^ The connection between the loop region and a short helix within ECL is further stabilized by a coordinated Na^+^ ion.^4^

The disulphide bond between Cys 114 and Cys 199 is defined in our cryo-EM density map. In the region of the disulphide bond between Cys 192 and Cys 198, on the other hand, the cryo-EM density is difficult to interpret, and the Na^+^ ion is not apparent in our structure.

## Discussion

The β1-adrenoceptor – AmpC β-lactamase fusion protein was designed to increase the mass of the receptor to improve the signal-to-noise ratio in the acquired cryo-EM data, and to introduce structural features that facilitate angular assignment of the particles in single-particle analysis, with the final aim of establishing a widely applicable engineering strategy for solving GPCR cryo-EM structures without forming active complexes with G-proteins, arrestin or artificial binders.

Many fusion proteins that have been used in GPCR crystallography were connected to the receptor via loops. To achieve rigidity using loops, optimization of the linkers is typically required (e.g. Refs^28,29^). Fusion protein connections that work for crystallization may not necessarily be rigid in solution or even in crystals. For example, in a crystal structure of a nanobody-stabilized β^2^AR construct containing a T4 lysozyme (T4L) fusion protein, the connections between the receptor and the fusion protein were flexible, resulting in averaging out of T4L in the structure, producing no interpretable electron density for T4L.^30^ Other connections formed chimeric extended helices that have the potential to be rigid also in solution, outside the crystal lattice (e.g. Ref.^31^).

GPCRs can correctly fold even if the folding flow is interrupted by fusion proteins in intracellular loops. This tolerance towards insertions may be related to the presence of independent folding units. It has been shown in GPCR research that rhodopsin, but also other GPCRs, can be produced as separate fragments, and that these fragments can combine with matching counterparts to form receptors that are functional in receptor-binding.^28,32–34^

Here, we specifically searched for a fusion protein that is larger than thermostabilized apocytochrome b_562_ RIL (BRIL)(e.g. Ref.^31^) but can also be genetically inserted at the location of ICL3 to form rigid, extended helices with TM5 and TM6 of the GPCR.^15^

The resulting cryo-EM map of cyanopindolol-bound ultra-thermostable β1-adrenergic receptor comprises information about the overall architecture of the GPCR, including the seven-helix transmembrane core, helix 8, and the ligand binding mode.

The overall architecture of GPCRs comprises an extracellular part composed of the N-terminus and three extracellular loops, a seven α-helix ligand-binding and information-transducing transmembrane domain core, and an intracellular part including three intracellular loops, the amphipathic helix 8, and the C-terminus.^35^

Thermostability-based ligand binding assays showed that the receptor, in the context of the fusion protein, efficiently binds cyanopindolol (and propranolol), confirming structural integrity.

3D variability analysis of the cryo-EM map revealed conformational heterogeneities within the receptor and between the receptor and the fusion protein. Unlike the crystal structure, the cryo-EM map of the flexible protein complex reflects the dynamics, at the expense of precise side chain positions, which could only be determined for a subset of residues (mostly with aromatic side chains) in the structure. On the other hand, density for the ligand cyanopindolol, which binds deeply in the center of the extracellular region of the receptor, is clearly defined in the cryo-EM map and allows determination of the overall ligand-binding mode. Large scale conformational changes between transmembrane helices 5 and 6 are prevented by the helix connections to the fusion protein.

Helix 8 (H8) of β1AR is critical for receptor stability and activation,^36^ and open questions remain about specific helix 8 functions. The base of helix 8 is wedged between helices 1 and 7, and appears like a lever arm, similar to a single α-helical domain.^37^ It seems plausible that this placement of H8 may stabilize the helix 1-7 core transmembrane bundle, and that leverage from H8 could potentially affect the conformation of the core and have a function in mechanosensing. Involvement in such a function of helix 8 has been suggested for GPCRs which are known to be mechanosensitive.^38^

In a physiological environment, helix 8 is anchored to the cytoplasmic face of the cell membrane through palmitoyl groups.^24^ In our construct sequence, the palmitoylation site (C358A) was removed by mutagenesis, and the protein was solubilized in detergent, hence no cellular membrane but only a micelle was present. Despite the absence of membrane anchoring or crystal contacts, helix 8 is defined in the cryo-EM map, in a typical position slanted upwards at a shallow angle towards the expected location of the cell membrane. Based on the localization of helix 8 relative to the lower rim of the DDM micelle, it is likely that the micelle stabilizes the helix through hydrophobic interactions.

The development of protein engineering approaches to enable the precise alignment of GPCR single particles in the absence of activating G proteins or arrestins is an active field of research. There have been recent reports of other strategies with similar aims, in most cases involving binding proteins such as nanobodies. For example, a newly developed nanobody that binds to an intracellular GPCR loop requires mutation of only a short stretch in the amino acid sequence of different receptors for binding.^39^ Furthermore, Mukherjee et al. generated synthetic antibodies that recognize apocytochrome b562 (BRIL), a frequently used fusion protein in GPCR structural biology, for cryo-EM applications.^40^ Zhang et al. explored fusion protein connections with one or two continuous single helices, often in combination with a nanobody,^41^ indicating that for structure elucidation at high resolution, either two stable helix connections, or one stable helix connection in combination with an additional, optimized linker or directed domain-to-domain contacts, are typically required.^41^ Xu et al. designed an approach comprising a third receptor-to-fusion protein connection by fusing one chain of the calcineurin heterodimer into ICL3, and the other chain to the C-terminus of TM 7 of the receptor.^42^ A similar three-point fusion using a GFP in combination with a GFP-binding nanobody has also been described for a transporter.^43^

In comparison to these and similar approaches, the AmpC β-lactamase we explored here is relatively large (much larger than for example BRIL), contributing strongly to a good signal-to-noise ratio even without binders, and makes no contacts to the receptor except for the direct helix fusions, allowing for the highest possible conformational freedom between receptor helices, except between TM5 and TM6, where movement is locked through the helix fusions. AmpC β-lactamase has an ideal geometry for insertion into the third intracellular loop of typical class A GPCRs and is a promising tool for the elucidation of GPCR structures in the absence of natural or artificial binding proteins. Because fusion proteins do not require complex formation, this approach holds great promise as a tool for the elucidation of cryo-EM structures with different ligands at an increased throughput for drug discovery research. Other potential applications include the elucidation of unliganded (apo) cryo-EM structures, and the elucidation of cryo-EM structures of GPCRs with low expression levels.

Our approach demonstrates the feasibility of creating a single construct where the fusion partner AmpC β-lactamase is genetically fused to transmembrane helices 5 and 6 (TM5 and TM6) of the receptor, eliminating the need for additional binding partners such as nanobodies. Further studies may confirm that this strategy can be widely applied to engineer other class A GPCRs similarly, thereby increasing the number of structures available for future research.

## Acknowledgements

We thank Mohamed Chami, Lubomir Kovacik and Carola Alampi at C-CINA Basel for support in cryo-EM data collection, the EM Facility at the PSI for technical support, and Takashi Ishikawa, Gebhard F.X. Schertler, Jan Pieter Abrahams, G.V. Shivashankar, Michel Steinmetz and Dmitry Veprintsev for supporting the project. We thank Hans Widmer, Sandra Jacob, Christian Wiesmann, Andreas Schenk and Gregor Cicchetti for interesting discussions about the project, Mara Wieser for support in protein production, Xavier Deupi for GPCR-related advice and Greta Assmann for computational support. We thank Daniel Böhringer for the coordination of the collaboration with the ScopeM Cryo-EM Knowledge Hub, and Julius Rabl for discussions and advice on the model building. This work was supported by grants from Novartis FreeNovation, the Promedica Stiftung Chur (1401/M) and the Swiss National Science Foundation (SNF SPARK, CRSK-3_190414) to R.M.B., and by grant 192760 from the Swiss National Science Foundation.

## Author contributions

The project was initiated and coordinated by R.M.B. The fusion proteins were designed by R.M.B.. The script for parsing the PDB was written by A.L. and S.B.. The protein was expressed by G.C. and T.B and purified by G.C.. Biophysical characterization was performed by G.C.. Grids for cryo-EM data collection were prepared and screened by G.C., I.M. and E.P.. Initial data analysis was performed by G.C. and E.P.. The cryo-EM structure was elucidated by P.A., G.C., I.M and R.M.B.. The final structure was analyzed and interpreted by R.M.B., P.A., I.M. and G.C.. The manuscript was written by R.M.B., G.C. and P.A., with important contributions from all authors.

## Materials and Methods

### Receptor and fusion protein selection

For the design of the chimeric protein, we used a β1AR construct with established expression and purification protocols. The sequence was based on the ultra-thermostable mutant β1AR-JM50,^17^ which contains the same mutations and truncations as the previous thermostabilized mutant β36-m23,^5^ but also comprises three additional thermostabilizing mutations: I129V, D322K, Y343L. The mutations from the β36-m23 construct served the following purposes: R68S, M90V, Y227A, A282L, F327A, F338M contributed to the increased thermostability and to the antagonist state,^4,44^ C358A eliminated the palmitoylation site, C116L resulted in improved expression levels, and the truncations at the N-terminus, C-terminus (and at intracellular loop 3) reduced flexible regions of the protein.^4,5,18^ The construct furthermore contains a D200E mutation, located in extracellular loop 2.

Fusion proteins were identified using a Java program. BioJava^45^ was used for the analysis. All structures in the Protein Data Bank as of February 2017 were considered. Secondary structure was assigned using the DSSP algorithm,^46^ implemented in BioJava. Candidates were considered if they had 15 residues annotated as helical near both termini. The terminal helices were required to fall within 11±1Å apart, as measured at the helix centroid. Finally, helices were required to be antiparallel within 15°. Structures which matched the filter criteria were then manually inspected, with 1FCO chosen as the most promising candidate.

The source code and documentation to run the program can be found in the following open repository: https://github.com/lafita/motif-search

### Transfection and stable cell line generation

HEK293S GnTI(-) cells^20^ were grown to ∼90% confluence in a cell culture dish at 37 °C, 5% CO_2_. Transfection was performed using Lipofectamine 3000 reagent (Invitrogen) according to the standard protocol. 100 µg/ml Zeocin were used for stable cell line selection.^19^

### Protein expression

The stably transfected cells were first upscaled in adherent cultures and then transferred to suspension cultures. Suspension cultures were grown in PEM (Protein Expression Medium, Gibco) supplemented with 10% FBS, 1% GlutaMAX (GIBCO), Zeocin (Invivogen) to a final concentration of 100 µg/ml and Penicillin-Streptomycin (PAN Biotech) to a final concentration of 100 U/ml. The suspension cultures were grown in a shaker-incubator at 120 rpm, 37 °C, 5% CO_2_. Cell densities were maintained between 800’000 and 1’500’000 cells/ml. Once the desired amount of cell culture volume was reached, the overexpression of the fusion protein was induced through the addition of tetracycline to a final concentration of 3 µg/ml. The induced cells were incubated for another 72 hours and then collected by centrifugation at 4’000 rpm, 15 min. at 4 °C and stored at −80 °C until further use.

### Membrane preparation and protein solubilization

All subsequent steps were performed at 4 °C or on ice. The thawed cell pellets were resuspended in a pellet-to-buffer ratio of 1:5 in 50 mM HEPES, pH 7.5, 1 mM MgCl_2_, 150 mM NaCl, supplemented with DNAse I and complete protease inhibitor cocktail (Roche). Cell membranes were disrupted using a dounce homogenizer. The homogenized material was ultracentrifuged at 230’000 rcf for 45 min., 4 °C. The supernatant was discarded. The membranes were gently resuspended using an ultra-turrax dispersing machine (IKA, USA) in solubilization buffer (50 mM HEPES, pH 7.5, 150 mM NaCl), supplemented with complete protease inhibitor cocktail (Roche). For solubilization, β-DDM was added to a final concentration of 1% (w/v). The mixture was stirred at 150 rpm for 2 hours at 4 °C. Another ultracentrifugation step was carried out to clarify the solution containing the solubilized receptor.

### Protein purification

The solubilized protein was incubated for 1 hour with washed and pre-equilibrated Strep-Tactin Superflow plus resin (Qiagen). The resin was then loaded into an Econo-Column Chromatography Column (Biorad) for gravity flow purification. The resin was washed 10 x with 2 volumes of wash buffer (50 mM HEPES, pH 7.5, 150 mM NaCl, 0.03 % (w/v) DDM). Stepwise protein elution was performed in wash buffer supplemented with 5 mM desthiobiotin (IBA-Lifesciences). The eluted protein was then treated with His-tagged human rhinovirus 3C protease (HRV 3C) to cleave off the C-terminal eGFP fusion and the Strep-tag. The protease was removed by Ni-NTA IMAC. The protein was then concentrated to 0.5 ml using a 50 kDa cutoff concentrator (Vivaspin, Sartorius, MWCO 50 kDa).

Size-exclusion chromatography was carried out on a Superdex 200 increase 10/300 GL column (GE Healthcare) in 50 mM HEPES, pH 7.5, 100 mM NaCl, 0.03% (w/v) DDM, 1 μM cyanopindolol. The protein was concentrated to 3 mg/ml using a 50 kDa cutoff concentrator (Vivaspin, Sartorius). The purified protein was frozen in liquid nitrogen and stored at −80 °C.

### Cryo-EM specimen preparation and data collection

For the final data collection, 3.5 μl of protein solution were applied onto the glow discharged Quantifoil Cu R1.2/1.3 200 grids (Electron Microscopy Sciences). The grids were blotted for 3s and plunge frozen into liquid ethane using a Vitrobot (Thermo Fisher Scientific) that was operated at 10 °C and 100% relative humidity. The data were collected on a 300 KeV FEI Titan Krios transmission electron microscope with a GATAN post-column Quantum-LS energy filter (20 eV zero-loss filter). 8,009 movies (40 frames each) were acquired in counting mode using SerialEM^47^ with a total dose of 64 e-/Å^2^ and pixel size 0.82 Å/pix. Data screening and selection of good micrographs were performed using in house scripts.^48^ Gain reference was estimated *a posteriori*^49^ using the *relion_estimate_gain* tool of the Relion5 software.^50^

### Single-particle analysis

Single-particle analysis was performed mostly in CryoSPARC v4.4.1 (Ref.^51^), however, many steps of the data processing were validated and optimized using Relion^50^ and cisTEM^52^ software packages. The processing pipeline is summarized in Supplementary Figure 1. Blob picking was initially used for the reference-free picking of 2,729,813 particles, out of which over a few iterations of 2D-classification 423,253 were selected and subjected for an *ab initio* initial model building based on randomly selected 20,000 particles. The particle set was then subjected to iterative rounds of 2D-classification and hetero-refinement for reducing the number of “bad” particles. Out of 164,992 “good” particles, 2D class-averages were obtained and were used as references for particle picking. The picked particles were combined again with those from the previous iteration of blob-picking, the duplicates were removed, and a stack of 2,865,326 particles was again subjected to iterative rounds of 2D-classification and hetero-refinements, resulting in the end in a stack of 183,537 “good” particles, which was further subjected to 3D-refinements using (Homo-/NU- and Local-refinement jobs). Various parameters and types of iterative jobs were tested to find an optimal workflow (including different manually created and dynamic masks, resolution limits and strength of low-pass filters), which resulted in a 3D-reconstruction at 4.3 Å resolution. Further 3D-classification jobs were performed to identify the largest 3D class-average out of 37,653 particles, reconstructed at 4.2Å resolution. The resulting 3D class-averages were analyzed to identify flexibility of the complex.

### Model building

Initial models of the fusion protein were generated using AlphaFold 2 and 3^25,53^, to include the overall architecture of the complete fusion protein in a single chain. Only AlphaFold2 predicted the geometry between the GPCR and the fusion protein as it was observed in the cryo-EM density. The model from AlphaFold2 was rigid body fitted into the cryo-EM map.

After initial real-space refinement, the GPCR part of the AlphaFold2 model was replaced by the model of a superposed high-resolution crystal structure^18^ (PDB:4BVN), to obtain a more precise starting model. After additional real-space refinement, the AmpC β-lactamase part of the AlphaFold2 model was in the same way replaced by a superposed high-resolution structure of this protein^54^ (PDB:6T3D) to create the model for further refinement. The overall structure of the complete fusion protein could be built into the cryo-EM map, from Serine 37 to Arginine 361 of the receptor. The model was refined in iterative steps of model building in Coot^55^ version 0.9.8.3 and real space refinement in phenix^56^ version 1.20-4459, using PDB:4BVN and PDB:6T3D for reference model restraints. MolProbity^57^ was used for validation of the geometry and stereochemistry of the model (Supplementary Table 1).

The model was refined as a continuously numbered single chain. After refinement, the GPCR part was renumbered to match the correct receptor numbering also in TM6 and 7. AmpC β-lactamase was renumbered to match the numbering in the crystal structure originally used in the construct design^16^ (PDB:1FCO) plus 1000, to keep it separate from the receptor numbering.

