## Supplementary Information for "Cryo-EM structure of a single-chain β1-adrenoceptor – AmpC β-lactamase fusion protein"

### Supplementary Figure

**Supplementary Figure 1)** Cryo-EM processing scheme

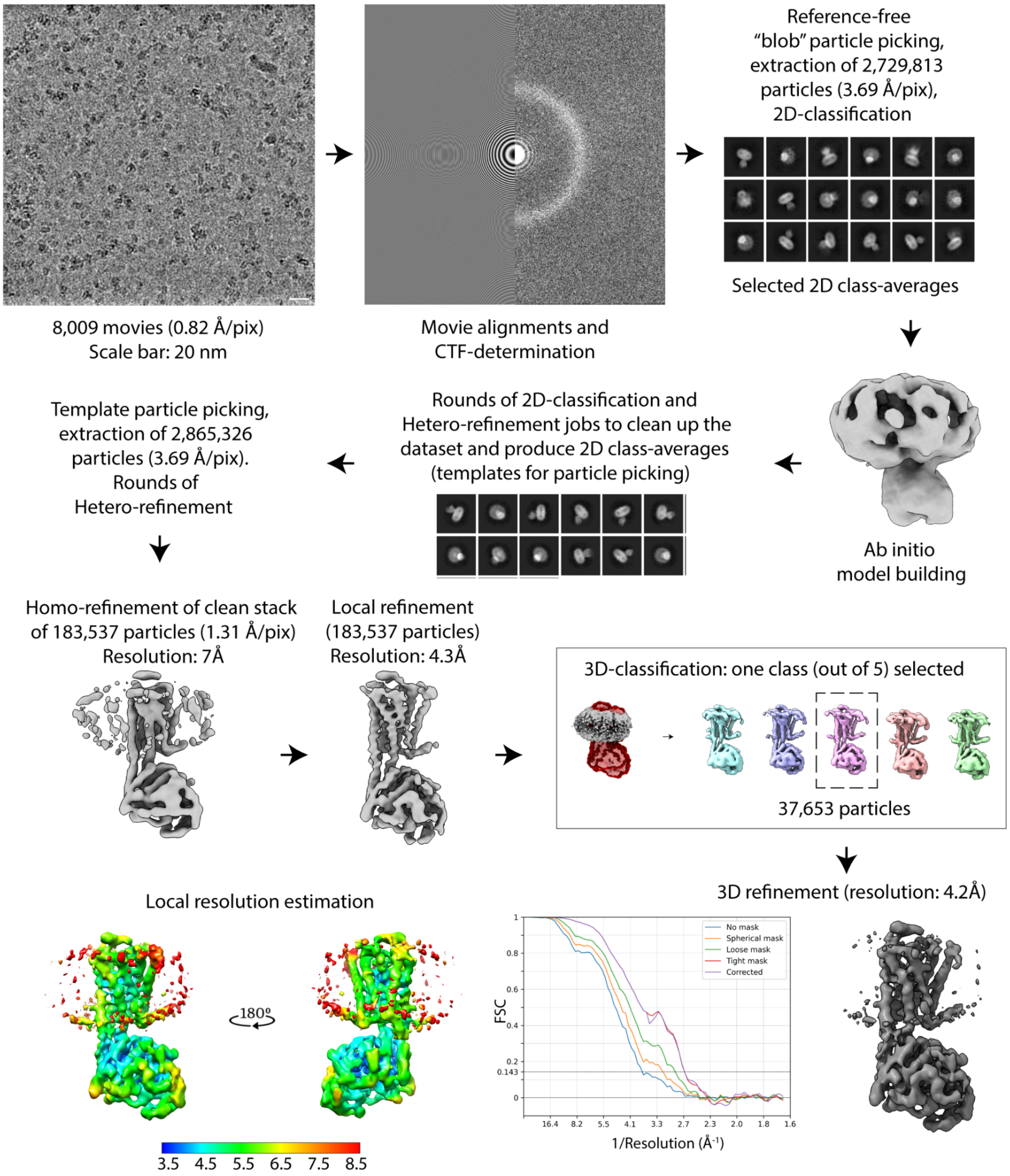

### Supplementary Movie

**Supplementary Movie 1)** 3D variability analysis demonstrates the first two variability components of the complex from three orthogonal side views.

### Supplementary Table

**Supplementary Table 1) Summary of MolProbity[57] statistics**

| | | | |
|---|---|---|---|
| **All-atom contacts** | Clashscore, all atoms | 16.71 | |
| | | **Raw count** | **Percentage** |
| **Protein geometry** | Poor rotamers | 3 | 0.60% |
| | Favored rotamers | 326 | 65.59% |
| | Ramachandran outliers | 0 | 0.00% |
| | Ramachandran favored | 613 | 97.30% |
| | Rama distribution Z-score | 0.48 ± 0.35 | |
| | MolProbity score | 1.86 | |
| | Cβ deviations >0.25Å | 0 | 0.00% |
| | Bad bonds: | 0 / 5029 | 0.00% |
| | Bad angles: | 0 / 6879 | 0.00% |
| **Peptide omegas** | Cis prolines: | 2 / 36 | 5.56% |
| **Low resolution criteria** | CaBLAM outliers | 14 | 2.2% |
| | CA geometry outliers | 4 | 0.64% |
| **Additional validations** | Chiral volume outliers | 0 / 793 | |
